# An interdisciplinary study around the reliquary of the late cardinal Jacques de Vitry

**DOI:** 10.1101/370908

**Authors:** Ronny Decorte, Caroline Polet, Mathieu Boudin, Françoise Tilquin, Jean-Yves Matroule, Marc Dieu, Catherine Charles, Aurore Carlier, Fiona Lebecque, Olivier Deparis

## Abstract

The reliquary of Jacques de Vitry, a prominent clergyman and theologian in the early 13th century, has experienced several transfers over the last centuries, which seriously question the attribution of the remains to the late Cardinal. Uncertainty about the year of his birth poses an additional question regarding his age at death in 1240. The reliquary, located in the Saint Marie d’Oigines church, Belgium, was reopened in 2015 for an interdisciplinary study around his relics as well as the Treasure of Oignies, a remarkable cultural heritage notably built from Jacques de Vitry’s donation. Anthropological, isotopic and genetic analyses were performed independently on the remains found in the reliquary. Results of the analyses provided evidence that the likelihood that these remains are those of Jacques de Vitry is very high: the remains belong to the same human male individual and the historical tradition about his age is confirmed. In addition, a separate relic (left tibia) was analysed and found to match with the remains of the reliquary (right tibia). The unique Jacques de Vitry’s mitre, made of parchment, was sampled non-destructively and the extracted parchment collagen was analysed by a proteomic method in order to determine the animal species. The results showed that, surprisingly, not all parts of the mitre were made from the same species. All together, these findings are expected to fertilize knowledge carried by historical tradition around the relics of Jacques de Vitry and his related cultural heritage.

## Introduction

Jacques de Vitry was a prominent clergyman and theologian, successively regular canon, bishop and cardinal of the Roman Catholic Church, who was active in Europe and Middle East during the first part of the thirteen-century (S1A Fig). His life and personality are mainly known from his writings (e.g. Historia Orientalis), crusade preaches and sermons. Unfortunately, no autobiography is available and facts about his youth are scarce. His date of birth is uncertain with two hypotheses coexisting: 1165-1170 (anonymous source, ca. 1250 [1]) or 1175-1180 (contemporary scholar Jean Donnadieu [2]), the former source being questionable (Jacques de Vitry neither studied theology in Paris in 1187 nor was the confessor of the king of France, as reported by the anonymous source). The date and place of his death, on the other hand, are known precisely as 1st May 1240 in Rome [3].

Having studied theology in Paris (between 1190 and 1208), he received episcopal consecration around 1210. He left Paris around 1208 to join the priory Saint Nicolas d’Oignies (funded in 1187) of the Diocese of Liège as an Augustinian canon regular. There, he met Marie d’Oignies (S1B Fig) and became the confessor and the biographer of the visionary, ecstatic beguine who died in 1213 at the age of 36 and was subject to popular devotions [4, 5]. Between 1212 and 1216, he preached the crusade against the Albigenses. Following his election as bishop of Saint John of Acre, he left the priory for the Holy Land in 1216. Following Damietta defeat in 1221, he decided to come back to Europe. A year after his return, in 1225, he resumed his itinerant life as preacher of the sixth crusade. In 1229, he was elevated to the College of Cardinals and settled in Rome where he died on May 1st 1240. His body was buried in the Dominican headquarter convent in Rome [6].

According to his testimonial will to lay at rest near to Marie d’Oignies, his remains were transferred to Oignies a year later and put in a marble monument, close to Marie’s one in the monastery church [7]. Several transfers of his relics took place in the next centuries. In 1636, the reliquary was open for transfer of his relics to a new place. Two teeth were removed on that occasion and given to clergymen for devotion [8]. On July 25th 1759, the reliquary was open for transfer to another place inside the church [9]. During the demolition of the priory in 1808, the relics were transferred to the Saint Martin church (Aiseau, Belgium) where they were put in a lead container, dated from 1844. After the collapse of the Saint Martin church building in 1970, the reliquary was opened in 1971 for the transfer of the remains to the Saint Marie d’Oignies church, Belgium, which was built in 1908 near the place of the ancient priory. The reliquary (and its content) is still displayed in this church today.

Throughout his life but also by testimony, Jacques de Vitry enriched the priory of Oignies with books, relics, tissues, vessels and other religious artworks [10]. He granted, among others, an ivory cross, a portable altar and two mitres, one of which being a unique object as it features miniatures on parchment. All these objects and others collected later constitute the Treasure of Oignies, a unique cultural heritage ensemble, which was recognized as such by the Belgian Federal State in June 2010 [11].

In 2015, the Archaeological Society of Namur (SAN), which has the scientific responsibility of the Treasure of Oignies, set up a consortium in partnership with several Belgian universities and research institutes in order to undertake an ambitious, interdisciplinary scientific study around the reliquary of Jacques de Vitry (referred as the CROMIOSS project). The study was motivated by the fact that the reliquary of the prominent clergyman and theologian has experienced several transfers during the last centuries, which seriously question the attribution of the remains to the late Cardinal. Uncertainty about the year of his birth poses an additional question regarding his age at death. Given the patrimonial importance of the Treasure of Oignies, the research consortium decided to englobe within its inquiry a material study of one of the bishop’s mitre (Fig 1), the only known example of a mitre composed of parchment. In spite of this exceptional feature, this mitre has so far received little attention from art historians.

**Fig 1.**
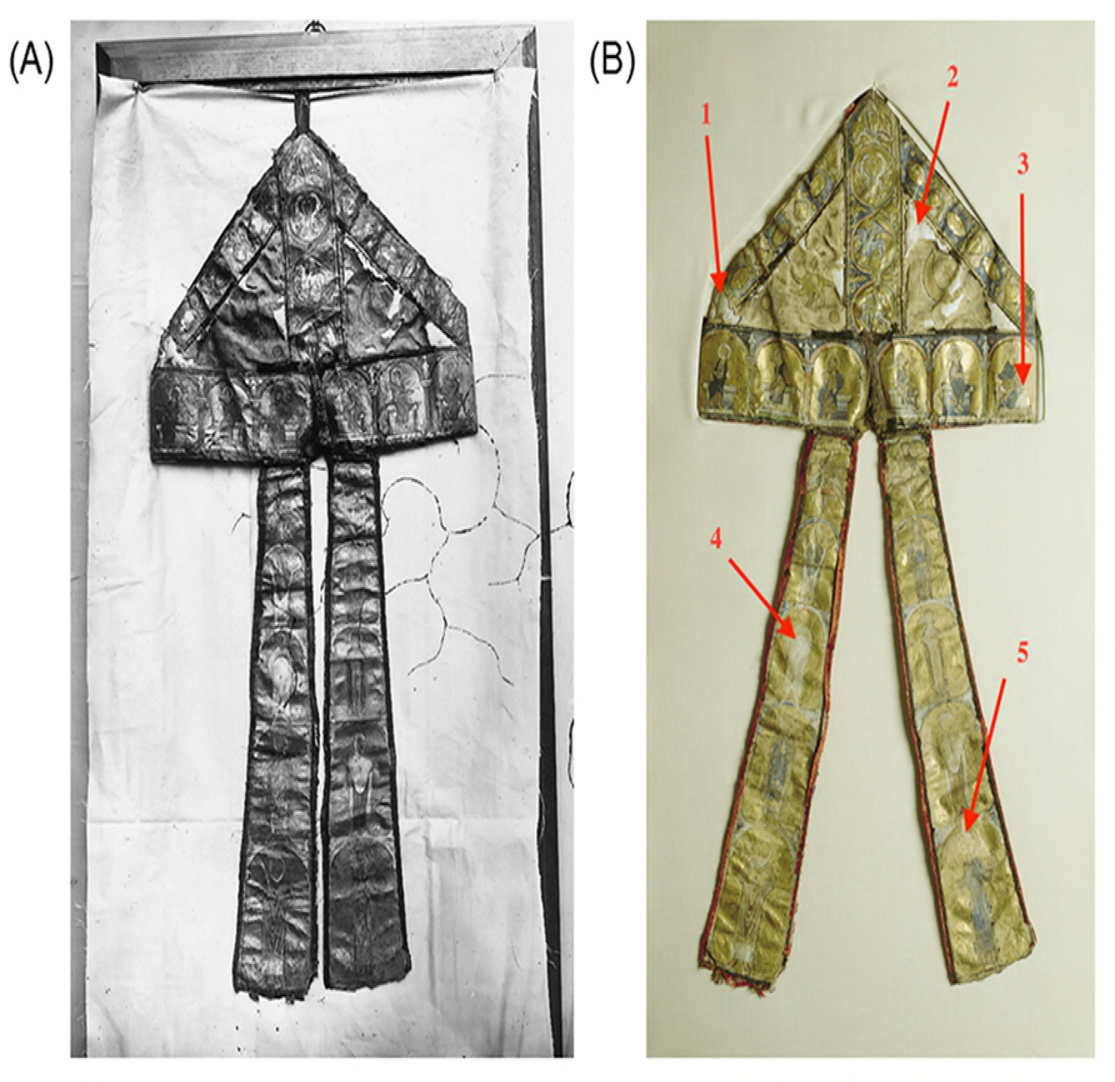
Jacques de Vitry’s mitre made of parchment. Donation of Jacques de Vitry to the priory of Oignies (collection of the Treasure of Oignies, Belgium). (A) Early photography (© Armand Dandoy, 1879). (B) Contemporary photography (© Vedrin, Guy Focant) with sampling spots indicated for parchment proteomic analyses (1, 2: cap; 3 cap border; 4, 5: left and right lappets).

Following the opening of the reliquary on September 8th 2015, human remains were found in the lead container. The interdisciplinary study reported hereafter relies on a critical confrontation between the historical tradition and the scientific results obtained from anthropological, isotopic, genetic and proteomic analyses.

## Materials and Methods

### Putative remains of Jacques de Vitry

With the approval of the competent authorities, the reliquary (S2A Fig) was opened on September 8th 2015, in the presence of the Bishop of the Diocese of Tournai, journalists and scientists. Caution was taken in order to avoid contamination of the relics by human DNA: restricted access, use of gloves and masks. Because the reliquary was previously open and relics were transferred several times since the second burial, it was first necessary to determine the human nature of the relics and to check whether or not they belonged to a single individual. A primary inspection of the lead container content was performed during the opening ceremony (S2B Fig). A wooden frame containing a tibia, which was supposed to belong to Jacques de Vitry, was also exhibited on that occasion. In fact, the story of this tibia is rather tumultuous. It was probably displayed in a private place and stolen at an unknown date. Found by the police in the 20^th^ century, it was first given back to the village of Oignies erroneously (actually, Oignies-en-Thiérache located in the Province of Namur, about 70 km from Oignies) before finally reaching the church Saint Marie of Oignies in the 20th century. In the reliquary, the remains were wrapped in a textile, which also contained small fragments of bone, plant remains, textile fibres, golden flakes and insect remains belonging to various groups of beetles (xylophagous (*Anobium punctatum*), detritivorous (*Ptinus* and *Tenebrio sp.*), necrophagous (*Trox scaber*) and granivorous (*Sitophilus granarius*)). The first three groups are part of taxa that are occasionally found in a funerary context and the presence of *Trox scaber* would suggest that organic material remnants, i.e. muscles, skin or hair, were still present. The reliquary, the frame and their contents were then transferred to the University of Namur (Laboratory of Anatomy) in order to take bone samples for further genetic and isotopic analyses.

### Anthropological study

A detailed inventory of the remains found in the reliquary was first established in order to estimate the minimum number of individuals. The sex determination of the studied individual was complicated by the absence of pelvic bones, which show the greatest sexual dimorphism. For this reason, we applied a sex determination method using cranium characteristics [12] and discriminant functions based on tibia measurements [13, p. 240-242]. In order to estimate the age at death, we used the dental wear [14], the closure of the cranial sutures [13, p. 120-121] and cementum analysis [15]. The stature was estimated using equations of Olivier et al., and computed from the length of long bones originating from a French sample [16].

### Isotopic study

Collagen extraction was performed following Longin’s (1971) method. A 1% NaOH wash step (15 minutes) was introduced between demineralization and hydrolization steps. First, all the bone samples were demineralized in 10 ml 8% HCl for 20 minutes, and rinsed with MilliQ^TM^-water. After that, each sample was immersed for 15 minutes in 1% NaOH, and again rinsed with MilliQ^TM^-water. Then, after adding 1% HCl for neutralization, it was washed with MilliQ^TM^-water. For all the steps mentioned above, Ezee-filters were used. Gelatinization of the extract was done in water (pH 3), at 90° C for 12 hours. The resulting gelatin was filtered with a Millipore 7 micrometre glass filter, and freeze-dried. Age determinations (^14^C analyses) were carried out on the AMS instrument at the Royal Institute for Cultural Heritage (KIRK-IRPA), Brussels (Lab code RICH-) [17]. CO_2_ was released by sample combustion in the presence of CuO and Ag. Graphitization was performed with H_2_ over a Fe catalyst. Targets were prepared at KIK-IRPA [18]. ^14^C calibrations were performed using OxCal (version 3.1) [19] and the IntCal13 calibration curve date [20]. The C:N ratio, δ^13^C and δ^15^N analyses were performed on a Thermo Flash EA/HT elemental analyser, coupled to a Thermo Delta V Advantage Isotope Ratio Mass Spectrometer via ConFlo IV interface (Thermo Fischer Scientific, Bremen, Germany). Standards used were IAEA-N1, IAEA-C6, and internally calibrated acetanilide. Analytical precision was 0.25‰ for both δ^13^C and δ^15^N based on multiple measurements of the standard acetanilide.

### Genetic study

DNA extraction and amplification were performed at UNamur and KULeuven in dedicated laboratories for ancient DNA analysis. Standard precautions for contamination were taken including separated areas for pre- and post-PCR procedures, restricted access to pre-PCR areas, multiple negative controls in the DNA extraction and amplification reactions, replication of DNA extractions and PCR reactions, contamination control with DNA profiles of all laboratory staff.

At UNamur laboratory, bone nuclear DNA was extracted according the procedure described by Mundorff and Davoren [21]. Briefly, the petrous bone and the tibias were cleaned with 10% bleach and bone powder was withdrawn by drilling with a decontaminated 8-mm drill under a previously decontaminated chemical hood. Nuclear DNA was then extracted from ≈3 g of bone powder using the Set Buffer Trace Bone kit and Nucleospin DNA Trace from forensic sample kit (Macherey-Nagel, Germany). Nine different microsatellite markers were amplified on the ancient bone DNA and contemporary DNA (used as control DNA) using the AmpFlSTR Minifiler PCR Amplification kit (Applied Biosystems) and were further analysed with GeneMapper on the Abi 3130xl Genetic Analyser (Applied Biosystems) at URBE (UNamur). Standard PCRs of 5 Y chromosome short tandem repeats (DYS19, DYS389II, DYS448, DYS456, DYS635) were performed using sets of primers designed by Kwon and coworkers [22].

At KULeuven laboratory, the procedures described by Ottoni et al. [23] were used for decontamination and grinding of the bone material and the two upper teeth samples, as well as for DNA extraction through silica-based spin columns (QIAquick PCR Purification Kit, Qiagen). One of the teeth was reused for genetic analysis after cement chrono-analysis. This tooth was embedded in epoxy resin and cut into two so that only the dentin and the root channel could be removed with a decontaminated dental drill. Multiplex DNA amplification of autosomal short tandem repeats (STRs) and STRs from the Y chromosome was done respectively according to Dognaux et al. and Larmuseau et al. [24–26] except for the number of cycles, which was raised to 34. The autosomal multiplex of 9 STRs (fragment size between 70 and 275 bp) includes also primers to amplify a sequence in an intron of the Amelogenin gene present on the sex chromosomes (123 bp on the X and 129 bp on the Y) [27]. Three multiplexes were used for the Y-chromosome STRs including in total 40 STRs with a size range between 74 and 420 bp. Fragment analysis was performed on an Applied Biosystems 3130XL Genetic Analyzer (Thermo Fisher Scientific) with data analysis using GeneMapper ID v3.2 software (Thermo Fisher Scientific). Analysis of the first and the second hypervariable segments (HV-I and HV-2) of the mtDNA control region was accomplished by amplification of, respectively, five and two overlapping fragments ranging in size from 109 to 166 bp, followed by direct sequence analysis according to Ottoni et al. [23]. Forward and reverse sequencing was performed using the BigDye Terminator v3.1 Cycle Sequencing Kit (Thermo Fisher Scientific) according to the protocol of the manufacturer. Sequence analysis was done on an Applied Biosystems 3130XL Genetic Analyzer (Thermo Fisher Scientific) with data analysis using DNA Sequence Analysis Software v5.2 (Thermo Fischer Scientific) and aligning of the sequences against the revised Cambridge Reference Sequence using BioEdit v7.0.4 [28, 29].

### Proteomic study

Non-invasive collagen extraction and sample preparation were done following the ZooMS method [30].

#### Sampling procedure

Sampling was performed in a clean area inside biological laboratory, with the mitre laid down on a clean table. Sampling consisted in gentle rubbing of the surface of parchment parts of the mitre (S3 Fig). For each sampling, a new piece of PVC erasers (Mars, Staedler) and new nitrile gloves were used, and the table was cleaned with isopropanol. Samples (eraser crumbs containing parchment collagen) were taken in duplicate at different locations where the parchment was bare (non-decorated parts): on the cap as well as on the right and left lappets (Fig 1B). Eraser crumbs were collected in a 1.5ml Eppendorf tube and were stored at 4°C until collagen extraction.

#### Collagen extraction and digestion

50 μl of NH4HCO3 50mM buffer was added to each sample. Eppendorf tubes were spin down at maximum speed in a centrifuge. 200 ng of trypsin (Promega) was added to each sample and incubated during 4 hours under light agitation. The samples were acidified with a solution of trifluoroacetic acid (TFA) to a final concentration of 1% (vol./vol.). Eppendorf tubes were centrifuged during 5 min at maximum speed and supernatant containing the peptides were transferred to another Eppendorf vial.

#### Peptides desalting and concentration

ZipTip C18 (Millipore) pipettes were used. After washing and conditioning of the ZipTip according the manufacturer’s instructions, the peptides were loaded and desalted with a solution of H_2_O, 0,1% TFA (vol./vol.). Peptides elution was done with 10 μl of 80% actonitrile (ACN) / 0.1% TFA (vol./vol.). All samples were then vacuum dried (Heto) and recovered with a solution of 2% ACN/ 0.1% TFA (vol./vol.).

#### Mass spectrometry analysis

All samples were analysed using liquid chromatography (UltiMate 3000, Thermo Systems) coupled to electrospray tandem mass spectrometry (MaXis Impact UHR-TOF, Bruker) (LC-MSMS). The digests were separated by reverse-phase liquid chromatography using a 75 μm X 150-mm reverse phase column (Acclaim PepMap 100 C18). Mobile phase A was 95 % H_2_O/5 % ACN, 0.1 % formic acid. Mobile phase B was 80% ACN/20% H_2_O, 0.1 % formic acid. The digest was injected, and the organic content of the mobile phase was increased linearly from 5 % B to 40 % B in 15 min and from 40 % B to 100 % B in 5min. The column effluent was connected to a Captive Spray (Bruker). In survey scans, MS spectra were acquired for 0.5 s in the m/z range between 50 and 2200. The 10 most intense peptides 2^+^ or 3^+^ ions were sequenced. The collision-induced dissociation (CID) energy was automatically set according to mass-to-charge (m/z) ratio and charge state of the precursor ion. MaXis and Thermo Systems instruments were piloted by Compass HyStar 3.2 (Bruker). Peak lists were created using DataAnalysis 4.0 (Bruker) and saved as mgf file for use with ProteinScape 3.1 (Bruker) with Mascot 2.4 as search engine (Matrix Science). Enzyme specificity was set to trypsin, and the maximum number of missed cleavages per peptide was set at one. Hydroxylation (KP) and oxidation (M) were allowed as variable modification. Mass tolerance for monoisotopic peptide window was 7 ppm and MS/MS tolerance window was set to 0.05Da. The peak lists were searched against a home-made collagen protein database and a contamination protein database. Scaffold software (Proteome Software) was used to validate protein and peptide identifications, and also to perform the search of species marker peptides. Our species marker database contained specific peptides that allowed us to differentiate *Capra hircus*, *Ovis aries* and *Bos taurus*.

## Results and discussion

### Results

#### Anthropological analysis of the remains

Inside the container, a cranium, 26 bones and 7 teeth were found, wrapped in a black satin shroud sealed with staples and a red wax seal. Mandible, ribs, pectoral and pelvic girdle bones were absent (Fig 2A), precluding efficient identification of the sex. No lower teeth were found and 9 upper teeth were missing: 7 were lost post mortem and 2 ante mortem.

**Fig 2.**
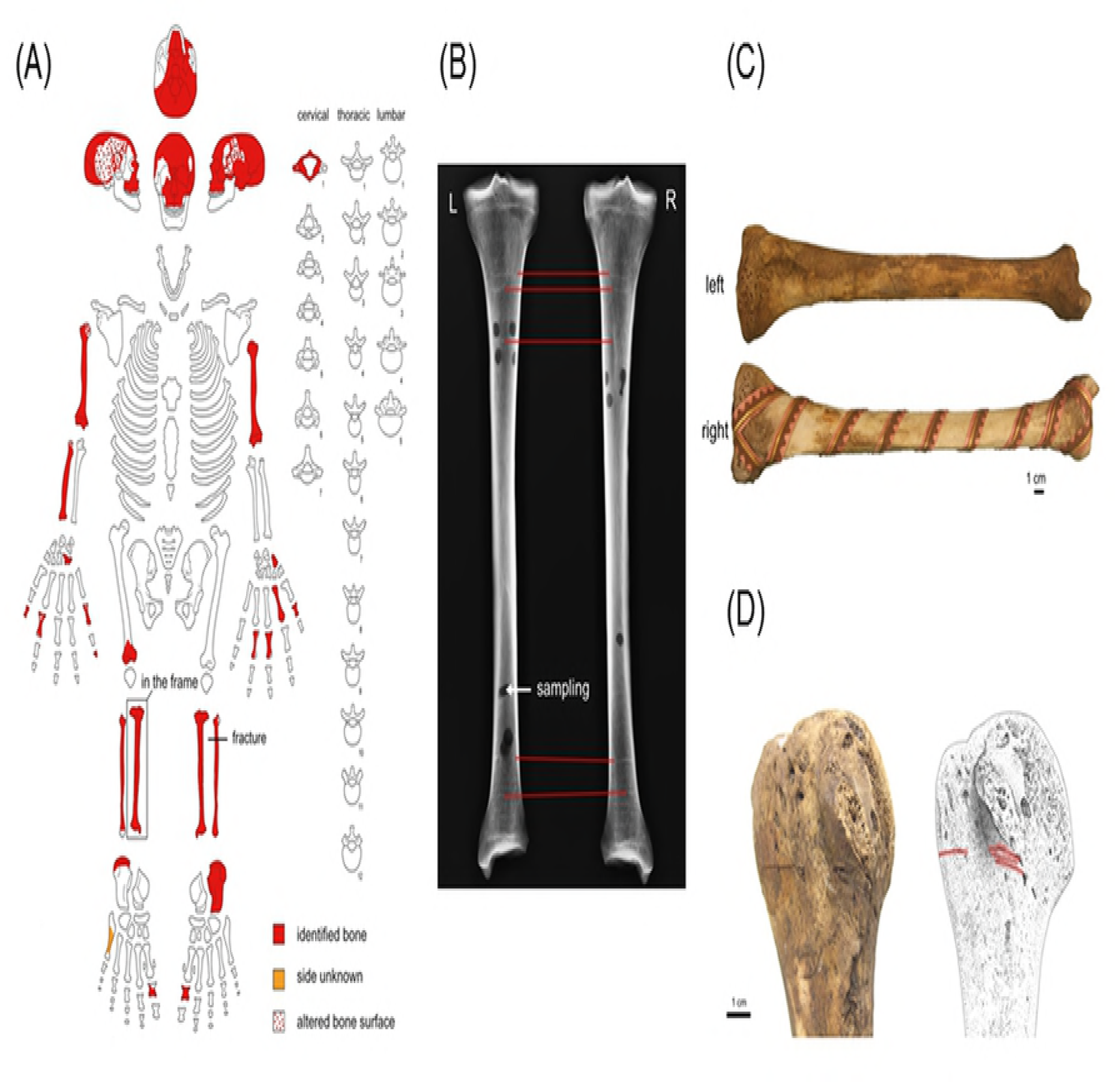
Remains found in the reliquary. (A) Inventory of the remains. (B) Radiography of the left and right tibias. (C) Left tibia (top) and right tibia (bottom). (D) Evidences of cuts on the bones (articulations).

Visual and metric inspection of the remains led us to conclude that they belong to the same individual. Metric comparison between the right and left tibias (Fig 2C) revealed almost no difference, except the discoloured aspect of the right tibia (the one put in the frame as a distinct relic), which might be the result of cleaning treatment applied before framing. Radiography of both tibias (Fig 2C) showed that the internal structures of both tibias exhibit strong similarity (concordance of Harris lines), which confirms they belong to the same individual.

Five cranial female characteristics (slightly delimited glabella, arched traces of nuchal lines and delimited supraorbital ridge; smooth external occipital protuberance and vertical frontal bone) and two male features (large mastoid process and quadrangular orbits) were identified. Inspection of tibias morphologies produced male or female diagnostics, depending on the morphological character considered.

Dental wear indicated an age ranging from 45 to 55. Cranial vault sutures indicated death between 30 and 60. Analysis of tooth cementum gave an age estimate from 53 to 62. The “consensus biological age” was around 55, which was supported by the fact that the specimen presented a few signs of degenerative osteoarthritis. Stature was estimated, based on the left tibia length. If the individual were a male (female), his (her) height would have been around 1.66 (1.62) meter.

Close visual inspection of the bones revealed peculiar, unexpected and interesting features: cut marks (Fig 2D) in the periphery and within articulations of the right shoulder, the right and left elbows, the right and left knees. These cuts were most likely made in order to remove muscles and ligaments.

#### Isotopic analysis of tibias and skull fragments

Isotopic analyses (δ^13^C, δ^15^N and ^14^C) were carried out on two samples taken from the left tibia, one sample from the right tibia and one sample from the skull. The results are shown in Table 1.

**Table 1.** Results of isotopic analyses: sample laboratory codes and names, ^14^C ages (BP), calibrated ages (2σ), stable isotope fractionations (δ^13^C and δ^15^N), C:N ratio of collagen extracted from different bone samples from the remains.

| Lab-code | Sample name | $^{14}\text{C}$ age (BP) | Calibrated age (2 $\sigma$) | $\delta^{13}\text{C}$ (‰) | $\delta^{15}\text{N}$ (‰) | Atomic C:N ratio |
| --- | --- | --- | --- | --- | --- | --- |
| RICH-22318.1.1 | Left Tibia 1 | 915±28 | 1030AD (95.4%)<br>1190AD | -19.9 | 10.8 | 3.3 |
| RICH-22318.2.2 | Left Tibia 2 | 903±28 | 1030AD (95.4%)<br>1210AD | -20.0 | 11.0 | 3.3 |
| RICH-22319.2.1 | Right Tibia | 896±26 | 1040AD (95.4%)<br>1220AD | -19.8 | 11.2 | 3.2 |
| RICH-23856 | Skull | 957±26 | 1020AD (95.4%)<br>1160AD | -19.0 | 11.0 | 2.9 |

BP, Before Present. $\sigma$ , standard deviation.

The C:N ratio of the collagen of all samples felt within the C:N range proposed by De Niro [31] for well-preserved collagen, namely between 2.9 and 3.6. The stable isotopes (δ^13^C, δ^15^N) results suggested that both tibias and the skull might belong to the same individual. The ^14^C results confirmed this hypothesis by passing positively the χ^2^-test. The average of the four ^14^C dates was calculated: 919±13BP (χ^2^-test: df=3, T=3.3(5% 7.8)), which led to a skeleton date between 1040 and 1170AD (95.4% probability) after calibration (Fig 3).

**Fig 3.**
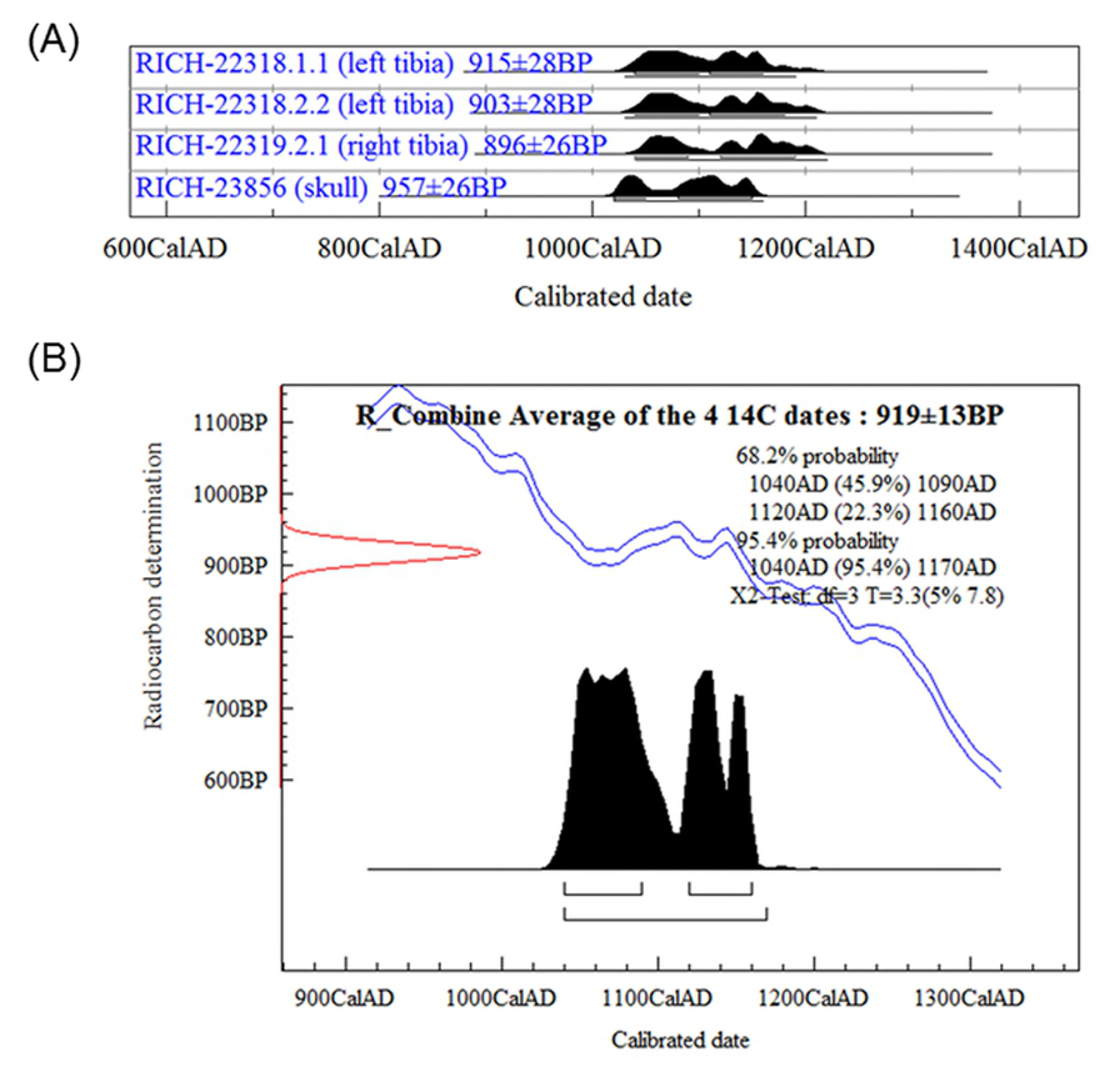
Radiocarbon dating results from the remains. (A) Calibrated ages (2σ) of the left tibia, right tibia and skull. (B) Average age of the four ^14^C dates determined for these remains.

#### Genetic analysis of the remains

Analysis of nuclear DNA from the petrous bone and both tibias was first performed at UNamur using 9 human DNA microsatellite markers. However, the analysis of DNA extracts from the remains turned out to be not reproducible among the different samples and the presence of the Y chromosome could not be established (data not shown). A standard PCR was then used to amplify 5 short tandem repeats (STR) of the Y chromosome using the same DNA samples. The Y chromosome was found in the petrous bone DNA, whereas the tibias DNA did not give reproducible results (Fig 4). These results suggest that the genetic material is strongly damaged in the tibias and, to a lesser extent, in the petrous bone.

**Fig 4.**
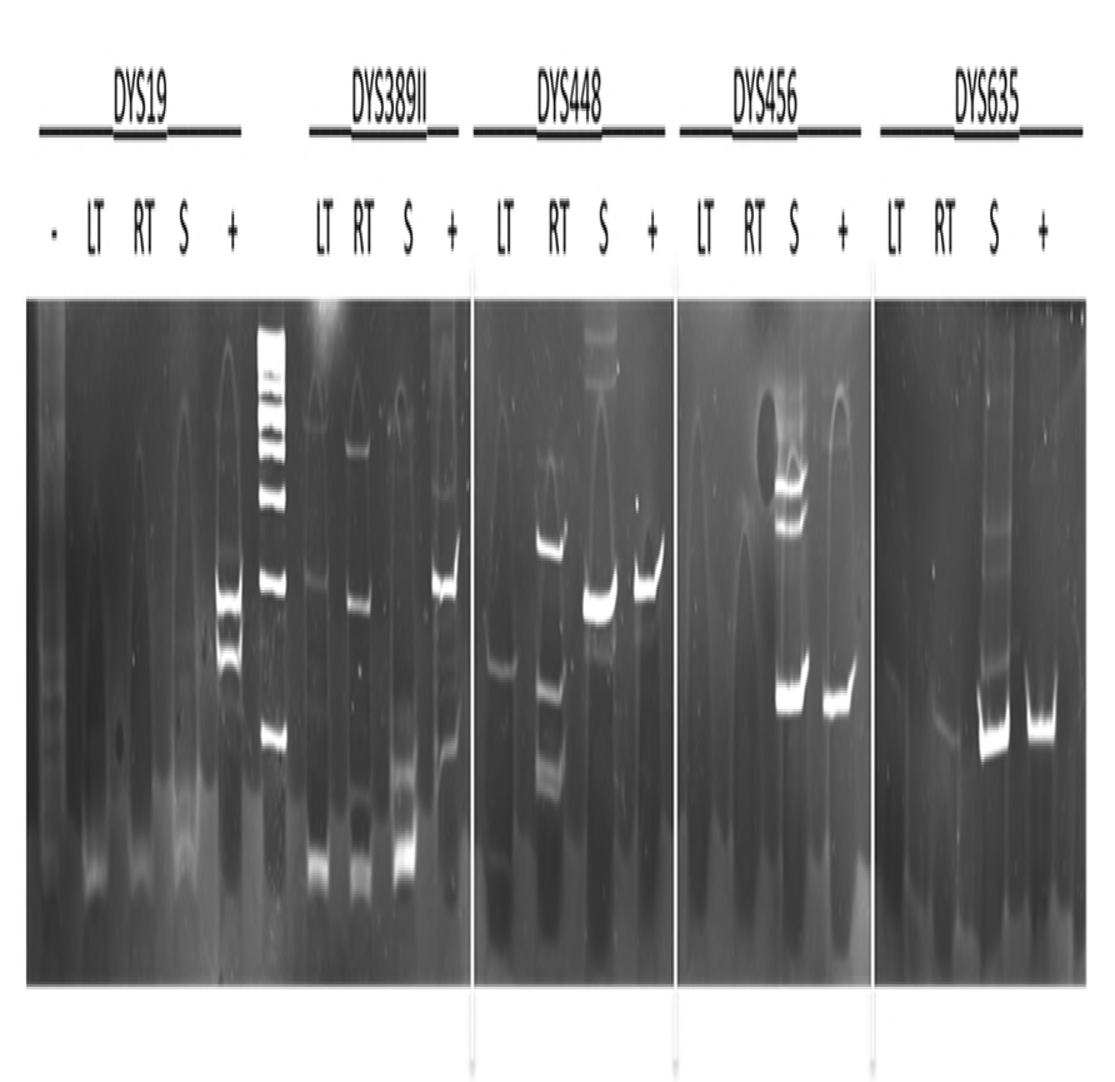
PCR amplification of 5 STRs from the Y chromosome. Left tibia (LT), right tibia (LT), petrous bone (S), negative control (-), positive control (+).

Genetic analysis of the remains was performed independently at KULeuven following standard ancient DNA (aDNA) protocols. Five samples were analysed: tibia powder, tooth embedded in epoxy resin (sample reused after cement chrono-analysis), a petrous bone fragment and two upper teeth. Analysis of short tandem repeats from the Y chromosome on the tibia DNA extract gave no results, indicating that the quantity and/or quality of the extracted nuclear DNA were insufficient or that the DNA originated from a female. Identical results were obtained for the resin embedded tooth and the petrous bone DNA extracts. The results obtained by UNamur and KULeuven for nuclear DNA are consistent with the treatment of the remains attributed to Jacques de Vitry (discussed below) which would have impacted the preservation of the DNA [32, 33]. In contrast, partial profiles for both autosomal (including sex determination with amelogenin gene) and Y-chromosome STRs were obtained with the two upper teeth DNA extracts (Table 2).

**Table 2.** Genotyping results of the analysis of nine autosomal STRs, Amelogenin and 40 Y-chromosome STRs in DNA extracts from two upper teeth.

| Autosomal STRs | AMEL<sup>1</sup> | D2S441 | D1S1656 | D12S391 | D10S1248 | D21S11 | D22S1045 | D18S51 | D1S1677 | FGA |
| --- | --- | --- | --- | --- | --- | --- | --- | --- | --- | --- |
| Amplicon length (bp) | 123-129 | 72-112 | 115-168 | 169-227 | 78-142 | 150-217 | 68-119 | 139-219 | 74-118 | 119-278 |
| Upper tooth 1 | Y | 11 | 16.3 | - | 14 | - | - | 14 | 13,15 | 18,(22)<br>[20] |
| Upper tooth 2 | Y | (11),14 | - | - | 14<br>[13] | 32.2 | - | 14,(20)<br>[19] | 13,(15) | 18,22 |
| Y-STRs | DYS438 | DYS392 | DYS458 | DYS385 | DYS460 | DYS481 |  |  |  |  |
| Amplicon length (bp) | 87-140 | 90-130 | 130-160 | 237,316 | 95-123 | 105-168 |  |  |  |  |
| Upper tooth 1 | 9 | 11 | 14 | 12,13 | 12,13 | 22 |  |  |  |  |
| Upper tooth 2 | - | - | 14 | - | 10 | - |  |  |  |  |

<sup>1</sup> Amelogenin.
[] PCR fragments attributed to stochastic effects of either high stutter or drop-in; () alleles with lower peak height (heterozygote imbalance).

The autosomal STR results were reproducible between the DNA extracts as well between different PCR reactions indicating that the two teeth originated from the same individual. The characteristics of these profiles (drop-out of alleles or loci, drop-in of one or two alleles, high stutter frequency, heterozygote imbalance) were consistent with ancient DNA, where DNA damage and fragmentation lead to non-amplification of alleles or loci, and where low amounts of DNA lead to stochastic effects (drop-out, high stutter effect and heterozygote imbalance) that will influence the results [34]. The fact that very partial profiles were obtained for the Y-chromosome STRs can be explained by the higher sensitivity of the autosomal STR multiplex for low amounts of degraded DNA, except for the Amelogenin amplicons where the X-allele could not be amplified in the DNA extracts of the two upper teeth.

All five DNA extracts were also subjected to the analysis of the non-coding region of human mtDNA. In contrast to the nuclear DNA results, all DNA extracts with the exception of the DNA extract from the tibia powder revealed a mitochondrial (mt) sequence (Table 3).

**Table 3.**
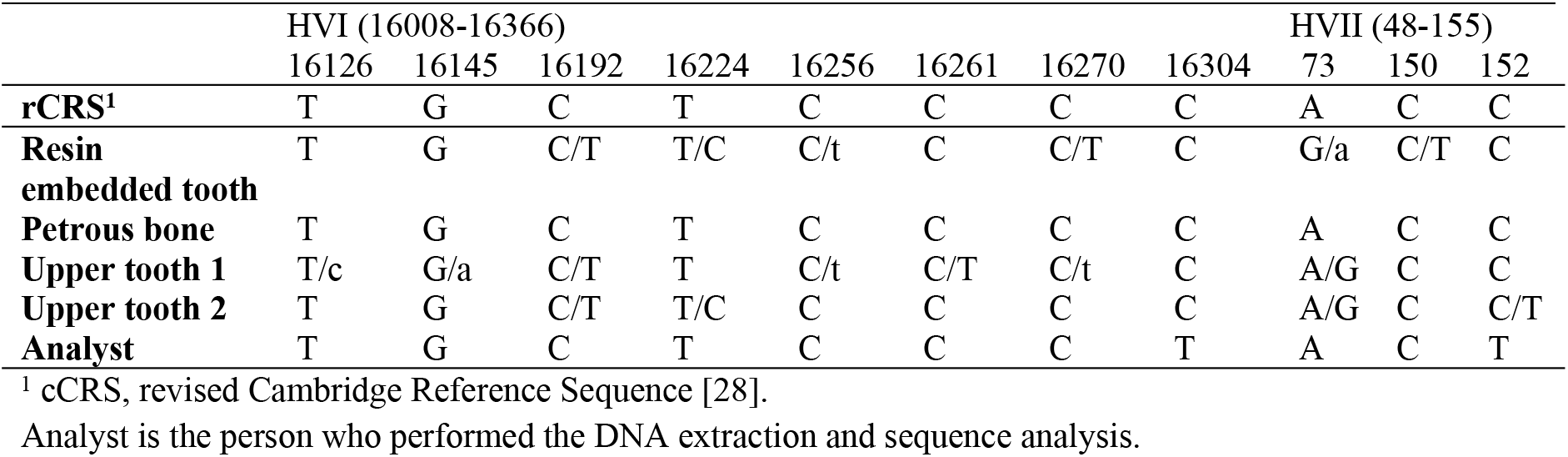
Results of mtDNA Sanger sequencing for different bone and teeth samples from the remains.

The samples of the reused tooth and the two upper tooth samples showed at several positions the presence of two nucleotides, which is evidence for the presence of exogenous DNA in the DNA extracts. The origin of this DNA remains unknown (laboratory contamination can be excluded) and can probably be related to the handling of the remains during the different historical transfers. The fact that four out of five DNA extracts showed preservation of human mtDNA can be explained by the higher copy number of mtDNA and, to a lesser extent, additional protection against degradation by the double membrane of the mitochondrion which would increase the chance to extract preserved mtDNA from archaeological remains [35, 36]. The mtDNA sequence of the DNA extract from the petrous bone fragment revealed no evidence for exogenous DNA and was identical to the revised Cambridge Reference Sequence [28]. This mtDNA sequence, the most frequent (about 10%) in West Eurasian populations (https://empop.online/;v3/R11), could also be present in the other DNA extracts, which would be consistent with the hypothesis that all the skeletal remains belong to the same individual. This hypothesis would also imply that the origins of the exogenous DNA in the other samples are different, which is supported by the different mtDNA sequences observed for the two upper teeth where reproducible nuclear DNA profiles were obtained.

#### Proteomic analysis of collagen from parchment parts of the mitre

Identification of the origin species of parchment parts of the mitre was carried out by proteomic analysis of the collagen protein extracted using a non-invasive sampling method (S3 Fig). Searches for species biomarkers gave unambiguous results on samples taken from four different locations on the mitre: cap, lower part of cap, left and right lappets (Fig 1B). Origin species (sheep, calf or goat) were identified in parchment samples according to the number of occurrences of species-specific peptide markers obtained during sequencing of collagen I and III proteins (Table 4).

**Table 4.** Results of species identification in parchment parts of the mitre.

| Sample | Location | Species-specific peptides (SSP) detected | Number of SSP validated and sequenced | Identified species |
| --- | --- | --- | --- | --- |
| 1 | Cap | 1 | 19 | *Ovis aries* (sheep) |
| 2 | Cap | 1 | 12 | *Ovis aries* (sheep) |
| 3 | Lower part of cap | 2 | 3 | *Bos Taurus* (calf) |
| 4 | Left lappet | 7 | 52 | *Bos Taurus* (calf) |
| 5 | Right lappet | 2 | 5 | *Bos Taurus* (calf) |

## Discussion

### History of the remains

In order to ease interpretation of the results of the scientific studies, we develop here additional historical considerations apart from the biography of Jacques de Vitry given above. Before Jacques de Vitry settled in Oignies, the relatively recent and poor priory did not attract attention. No doubt the arrival of Jacques de Vitry changed the destiny of the priory, favouring its enrichment by the acquisition of relics [37, 38]. The meeting with Marie d’Oignies had important consequence for the destiny of the future bishop and cardinal since it was at the origin of his decision to be buried in Oignies. Actually, it was a current practice in the high society of that time to notify by testimony the will to be buried in homeland in case of death occurred abroad. Repatriation of dead human bodies from long distances had practical consequences in terms of treatments applied to the corpse in order to ensure hygienic and safe transfer. This implied dismembering of the corpse and use of recipes to treat the remains [39-41]. It is noteworthy that this practice was forbidden in 1299 by Pope Boniface VIII as it was regarded as an abominable custom for Christians. About 50 years before the ban, in due respect to his testimonial will, it is reasonable to assume that the dead body of Jacques de Vitry has experienced dismembering and subsequent treatment of the remains. It is also noteworthy that a left tibia put in a wooden frame was kept as a separate relic (Fig 2C). No clear explanation has been given so far about the origin of this relic, except that it is attributed to Jacques de Vitry thanks to a label present in the frame and bearing the following notice in French: “Ceci est un ossement du Vénérable Jacques de Vitry. Cardinal et Evêque d’Acre en Palestine. Mort à Rome le 30 Avril 1240. Transporté à son Monastère D’Oignies l’année suivante, d’où provident cette précieuse Relique” wich means: “This is a bone from the Respectable Jacques de Vitry. Cardinal and Bishop of Acre in Palestine. Died in Rome on May 1st, 1240. Transported to his Monastery of Oignies next year, from where this invaluable Relic comes.” Therefore, a question naturally arises regarding the belonging of this framed tibia to the remains found in the reliquary.

### Are the remains those of Jacques de Vitry?

In the absence of historical portrait or description, traits of Jacques de Vitry remain unknown. No information is available regarding his stature, physiognomy and health. Moreover, we face the case of a secondary burial, not the original one. His age at death is uncertain: between 60 and 75. Regarding the sex, it turned out that the morphology of the (incomplete) skeleton did not allow us to determine it. Therefore, we decided to resort to genetic analysis with the hope to be able to determine the sex from preserved DNA. Regarding the age, anthropological results (dental wear, cranial vault and tooth cementum) tend to support the contemporary hypothesis regarding the birth year of Jacques de Vitry, with a slightly lower age estimate. The biological profile (age, sex, stature) obtained from the anthropological study did not allow us to deliver a verdict about the attribution of the remains to Jacques de Vitry. Cut marks on bones, on the other hand, are compatible with contemporary medieval practices used for hygienic and safe repatriation of dead human bodies. Evidences of cuts on several bones and around joints suggest dismembering of the corpse [42-44] in order to facilitate post mortem transfer. This hypothesis is supported by historical reports of contemporary dismembering practices used for remote burial and agrees with the translation of Jacques de Vitry remains from Rome to Oignies, on a distance of more than 1400 km. According to historical recipes, dismembering was followed by boiling of bones in water, wine or vinegar [39-41]. Such treatments would have damaged seriously the genetic material in the bones [45, 46].

The results of isotopic analyses of the remains (Table 1) show that the left tibia, the right tibia and the skull are from the same individual. Taking into account the standard deviations on the ^14^C ages, there are no significant differences between the dates. The upper bound of average ^14^C dates (1170AD) is anterior (70 years offset) to the year of death of Jacques de Vitry (1240AD). On the other hand, the stable isotopes analyses indicate a mainly terrestrial diet. If this happened to be the case, the skeleton could not be attributed to Jacques de Vitry. However, the δ^15^N value is quite enriched, which may indicate a small amount of fish consumption since a carnivore has a δ^15^N value of +8‰ (S1 Table) [47]. Based on the depleted δ^13^C value, it can be assumed this comes from freshwater fish and not marine fish, which has a more enriched δ^13^C value (compare Table 1 and S1 Table). Freshwater fish can have reservoir ages ranging between several hundreds of years and two thousand years, as demonstrated in [48]. A minimal consumption of freshwater fish with a large reservoir age can explain the offset between the year of death of Jacques de Vitry and the obtained radiocarbon ages. If this reservoir effect is present, then the remains are likely to belong to Jacques de Vitry.

The objective of the genetic analysis was to determine the sex of the individual whose remains were found in the reliquary. Based on the DNA results for the petrous bone and two upper teeth (sex determination with amelogenin gene and analysis of Y-chromosomal STRs), we are able to conclude that they originate from a male individual.

### Jacques de Vitry’s mitre

Jacques de Vitry bequeathed several personal objects, which constituted, among others, the Treasure of Oignies. Among these, the mitre having miniatures on parchment is not only unique but also intriguing: questions about his usage, origin and fabrication have not been addressed so far. The results of proteomic analyses showed that parchment of the cap was made from sheep, except the lower part where calf was identified. On the other hand, parchment of both lappets was made of calf. This difference of origin species between different parchment parts of the mitre is puzzling and has not been reported before. Was this choice of different animal species intentional or not? If yes, was it motivated by differences in parchment quality, price or availability according to the species? All these questions certainly deserve further investigations from the point of view of the history of art.

## Conclusion

An interdisciplinary study was carried out around the reliquary of the late cardinal Jacques de Vitry, a prominent clergyman and theologian who was active in Europe and Middle East during the first part of the thirteen century. Results of anthropological, isotopic and genetic analyses provided evidence that the likelihood that the remains found in the reliquary are those of Jacques de Vitry is very high. Parchment parts of a mitre having belonged to Bishop Jacques de Vitry were analysed non-invasively by proteomic techniques and found to be made of different animal species. These findings are expected to fertilize knowledge carried by historical tradition around the relics of Jacques de Vitry and his related cultural heritage.

## Acknowledgments

The CROMIOSS project (www.lasan.be/la-recherche/projet-cromioss) was funded by a grant from the Fonds Jean-Jacques Comhaire of the Fondation Roi Baudouin (FRB), Belgium. Anne De Breuck (FRB) and Prof. Jean-Jacques Cassiman (KU Leuven, Belgium) are acknowledged for their support to the project. Jacques de Vitry’s mitre (donation from the Soeurs de Notre-Dame de Namur to FRB) belongs to FRB collections and is hold at Musée provincial des Arts anciens du Namurois – Trésor d’Oignies (TreM.a).

We thank Caroline Tilleux (Université Catholique de Louvain and Musées Royaux d’Art et d’Histoire, Belgium) for her help during examination of the remains. We also want to thank Jean-François Nisolle (Centre Hospitalier Universitaire Dinant Godine, Belgium) for radiographies of the tibiae, Benoît Bertrand (University of Lille, France) for age estimation using cementum analysis, and Anne-Marie Wittek (Association pour la Diffusion de l’Information Archéologique, Belgium) for drawing the sketch of bones with cut marks.

We thank Prof. Karine Van Doninck, Jonathan Marescaux and Catherine Demazy (URBE, University of Namur, Belgium) for giving access to the genetic analyser and for providing technical support.

Prof. Pierre Garin (Laboratory of Anatomy, University of Namur, Belgium) is acknowledged for hosting of the remains during the period of the project.

Finally, we are indebted to Jean Donnadieu (Université de Provence Aix-Marseille I, France), expert of Jacques de Vitry’s history.

## Author contributions

Conceived the study and requested authorizations for carrying analyses on Jacques de Vitry’s reliquary and mitre: FL AC. Genetic analyses: RD FT JYM. Anthropological analyses: CP. Isotopic analyses: MB. Proteomic analyses: MD. Mitre sampling: CC. Coordination of the writing of manuscript: OD. All authors discussed the results and approved the manuscript.

## Supporting information

**S1 Fig. Jacques de Vitry and Marie d’Oignies.** (A) Cardinal Jacques de Vitry on his deathbed (engraving). (B) Saint Marie d’Oignies (engraving).

**S2 Fig. Reliquary of Jacques de Vitry.** (A) The reliquary (© Vedrin, Guy Focant). (B) The remains found in the reliquary after opening on 8^th^ September 2015 (© Vedrin, Guy Focant).

**S3 Fig. Non-invasive sampling of parchment parts of Jacques de Vitry’s mitre.** Gentle rubbing of the parchment surface with a PVC eraser for proteomic analyses (© Vedrin, Guy Focant).

**S1 Table.** Isotopic fractionations of carbon (δ^13^C) and nitrogen (δ^15^N) according to diet.

| Bone collagen from animals having a 100% diet of | $\delta^{13}\text{C}$ (‰) | $\delta^{15}\text{N}$ (‰) |
| --- | --- | --- |
| C3-plants | -21 | +5 |
| Meat C3-herbivores | -18 | +8 |
| C-4 plants | -7 | +5 |
| Marine food | -13 | +18 |
| River fish | -24 | +16 |
| Lake fish | -20 | +16 |

Data reproduced from Lanting and van der Plicht (1996, 1998).

## References

[1] Huygens RBC, editor. Historia fundationis Venerabilis Ecclesiae Beati Nicolai oigniacensis ac Ancilla Christi Mariae Oigniacensis. Turnhout: Brepols Publishers; 2012. (Latin)

[2] Donnadieu J. Jacques de Vitry (1175/1180-1240). Entre l’Orient et l’Occident:l’Evêque aux trois visages. Turnhout: Brepols Publishers; 2014. (French)

[3] Auvray L, editor. Les registres de Grégoire IX, publiés et analysés d’après les manuscrits originaux du Vatican. Paris: Fontemoing; 1896. n° 5179. (Latin)

[4] Huygens RBC, editor. De Vitry J. Vita Mariae de Oegnies. Turnhout: Brepols Publishers; 2012. (Latin)

[5] Buisseret F. Histoire de la vie, miracles et translations de S. Marie d’Oignies. Louvain: Gerard Rivius; 1609. (French)

[6] De Cantimpré T. Vita Sanctae Lutgardis. Paris: Acta Sancotorum Juin/III; 1867, p. 257. (Latin)

[7] De Trois-Fontaines A. Chronica. Scheffer-Boischost Peditor. Leipzig; 1925, p. 950. (Latin)

[8] Moschus F. Caenobiarchia Ogniacensis, sive Antistitum Ogniacensium Catalogus; Auctore Francisco Moscho, Accessere Elenchus sacrarum reliquiarum, quae ibidem in cimeliarchiopie adservantur; et sanctorum vitae, qui ibidem qui escunt, donec illucescat dies a eternitatis in futurum. Omnia cura et labore Arnoldi Raissi. Douai: Bartholomé Bardou; 1636, p.p. 9–44. (Latin)

[9] Chartes du prieuré d’Oignies de l’ordre de Saint-Augustin. Poncelet E, editor. In: Annales de la Société archéologique de Namur. Namur: Société archéologique de Namur; 1912, pp. 31–101. (French)

[10] Rayssius A. Catalogus Sanctarius Reliquiarum. In: Moschus F. Caenobiarchia Ogniacensis, sive Antistitum Ogniacensium Catalogus; Auctore Francisco Moscho, Accessere Elenchus sacrarum reliquiarum, quae ibidem in cimeliarchiopie adservantur; et sanctorum vitae, qui ibidem qui escunt, donec illucescat dies a eternitatis in futurum. Omnia cura et labore Arnoldi Raissi. Douai: Bartholomé Bardou; 1636, pp. 48–53. (Latin)

[11] Actes de la journée d’étude Hugo d’Oignies. Contexte et perspectives. Namur: Société archéologique de Namur; 2012, p.58. (French)

[12] Ferembach D, Schwindezky I, Stoukal M. Recommendation for age and sex diagnoses of skeletons. J Hum Evol. 1980;9 517–549.

[13] Krogman WM, Işcan, M.Y. The Human Skeleton in forensic medicine, 2nd edition. Springfield: Thomas CC Publisher; 1986.

[14] Lovejoy CO. Dental wear in the Libben population: its functional pattern and role in the determination of adult skeletal age at death. Am J Phys Anthropol. 1985;68(1): 47–56.

[15] Colard T, Bertrand B, Naji S, Delannoy Y, Bécart A. Toward the adoption of cementochronology in forensic context. Int J Legal Med. 2015;129: 1–8.

[16] Olivier G, Aaron C, Fully G, Tissier G. New estimations of stature and cranial capacity in modern man. J Hum Evol. 1978;7(6): 513–518.

[17] Boudin M, Van Strydonck M, van den Brande T, Synal H-A, Wacker L. A new AMS facility at the Royal Institute for Cultural Heritage, Brussels, Belgium. Nuclear Instruments and Methods in Physics Research B. 2015;361: 120–123.

[18] Van Strydonck M, Van der Borg K. The construction of a preparation line for AMS- targets at the Royal Institute for Cultural Heritage, Brussels. Bulletin Koninklijk Instituut voor Kunstpatrimonium. 1990–1991;23:228–234.

[19] Bronk Ramsey C. Radiocarbon calibration and analysis of stratigraphy: the OxCal program. Radiocarbon 1995;37(2): 425–430.

[20] Reimer PJ, Bard E, Bayliss A, Beck JW, Blackwell PG, Bronk Ramsey C, et al. IntCal13 and Marine13 radiocarbon age calibration curves 0–50,000 years cal BP. Radiocarbon 2013;55(4): 1869–1887.

[21] Mundorff A, Davoren JM. Examination of DNA yield rates for different skeletal elements at increasing post mortem intervals. Forensic Science International: Genetics. 2017;8: 55–63.

[22] Kwon SY, Lee HY, Kim EH, Lee EY, Shin KJ. Investigation into the sequence structure of 23Y chromosomal STR loci using massively parallel sequencing. Forensic Science International: Genetics. 2016;25: 132–141.

[23] Ottoni C, Ricaut F-X, Vanderheyden N, Brucato N, Waelkens M, Decorte R. Mitochondrial analysis of a Byzantine population reveals the differential impact of multiple historical events in South Anatolia. Eur J Hum Genet. 2011;19: 571–576.

[24] Larmuseau MHD, Vanderheyden N, Jacobs M, Coomans M, Larno L, Decorte R. Micro- geographic distribution of Y-chromosomal variation in the central-western European region Brabant. Forensic Sci Int Genet. 2011;5: 95–99.

[25] Dognaux S, Larmuseau MHD, Jansen L, Heylen T, Vanderheyden N, Bekaert B, et al. Allele frequencies for the new European Standard Set (ESS) loci and D1S1677 in the Belgian population. Forensic Sci Int Genet. 2012;6: 75–77. doi:10.1016/j.fsigen.2011.05.003.

[26] Larmuseau MHD, Bekaert B, Baumers M, Wenseleers T, Deforce D, Borry P, et al. Biohistorical materials and contemporary privacy concerns-the forensic case of King Albert I. Forensic Sci Int Genet. 2016;24. doi:10.1016/j.fsigen.2016.07.008

[27] Sullivan KM, Mannucci A, Kimpton CP, Gill P. A rapid and quantitative DNA sex test: fluorescence-based PCR analysis of X-Y homologous gene amelogenin. Biotechniques. 1993;15: 636–8, 640–1.

[28] Andrews RM, Kubacka I, Chinnery PF, Lightowlers RN, Turnbull DM, Howell N. Reanalysis and revision of the Cambridge reference sequence for human mitochondrial DNA. Nat Genet. Nature Publishing Group; 1999;23: 147–147. doi:10.1038/13779.

[29] Hall TA. BioEdit: a user-friendly biological sequence alignment editor and analysis program for Windows 95/98/NT. Nucl Acids Symp Ser. 1999;41: 95–98.

[30] Fiddyment S, Holsinger B, Ruzzier C, Devine A, Binois A, Albarella U, et al. Animal origin of 13th-century uterine vellum revealed using noninvasive peptide fingerprinting. Proc Natl Acad Sci. 2015;112(49):15066–71. doi: 10.1073/pnas.1512264112. PubMed PMID: 26598667; PubMed Central PMCID: PMCPMC4679014.

[31] DeNiro MJ. Postmortem preservation and alteration of in vivo bone collagen isotope ratios in relation to palaeodietary reconstruction. Nature. 1985;317(6040): 806–809.

[32] Pruvost M, Schwarz R, Correia VB, Champlot S, Braguier S, Morel N, et al. Freshly excavated fossil bones are best for amplification of ancient DNA. Proc Natl Acad Sci U S A. National Academy of Sciences; 2007;104: 739–44. doi:10.1073/pnas.0610257104.

[33] Ottoni C, Koon HE, Collins MJ, Penkman KE, Rickards O, Craig OE. Preservation of ancient DNA in thermally damaged archaeological bone. Naturwissenschaften. 2009;96: 267–278.

[34] Ottoni C, Bekaert B, Decorte R. DNA degradation: Current knowledge and developments. In: Schotsmans EMJ, Márquez-Grant N, Forbes SL, editors. Taphonomy of human remains: forensic analysis of the dead and the depositional environment. John Wiley & Sons Ltd, Chichester, West Sussex; 2017. pp. 65–80.

[35] Schwarz C, Debruyne R, Kuch M, Mcnally E, Schwarcz H, Aubrey AD, et al. New insights from old bones: DNA preservation and degradation in permafrost preserved mammoth remains. Nucleic Acids Res. 2009;37: 3215–3229. doi:10.1093/nar/gkp159.

[36] Higgins D, Rohrlach AB, Kaidonis J, Townsend G, Austin JJ. Differential Nuclear and Mitochondrial DNA Preservation in Post-Mortem Teeth with Implications for Forensic and Ancient DNA Studies. PLoS One. 2015;10: 0126935. doi:10.1371/journal.pone.0126935.

[37] Huygens RBC, editor. De Vitry J. Vita Mariae de Oegnies. Turnhout: Brepols Publishers; 2012, p.144. (Latin)

[38] Wankenne A. La vie de Marie d’Oignies par Jacques de Vitry. Supplément par Thomas de Cantimpré. Namur; 1989, p. 169. (French)

[39] Paravicini-Bagliani A. Démembrement et intégrité du corps au XIIIe siècle. Terrain; 1992, p. 18, pp. 26–32. (French)

[40] Georges P. Mourir c’est pourrir un peu … Intentions et techniques contre la corruption des cadavres à la fin du Moyen Age. Micrologus, Nature, Sciences and Medieval Societies. 1999;VII: 359–382. (French)

[41] Weiss-Krejci E. Excarnation, evisceration, and exhumation in medieval and post-medieval Europe. In: Rakita GFM, Buikstra J, Beck L, Williams S, editors. Interacting with the dead. Perspectives on mortuary archaeology for the new millennium. Gainesville: University Press of Florida. 2005; pp. 155–172.

[42] Porta D, Amadasi A, Cappella A, Mazzarelli D, Magli F, Gibelli D, et al. Dismemberment and disarticulation: A forensic anthropological approach. J Forensic Leg Med. 2016;38: 50–57.

[43] Morcillo-Méndez MD, Campos IY. Dismemberment: Cause of death in the Colombian armed conflict. Torture. 2012;22 (Suppl 1): 5–13.

[44] Rutty GN, Hainsworth SV. The Dismembered Body. In: Rutty GN, editor. Essentials of Autopsy Practice: Innovations, Updates and Advances in Practice. London: Springer 2014; pp. 59–87.

[45] Arismendi JL, Baker, LE, Matteson KJ. Effects of Processing Techniques on the Forensic DNA Analysis of Human Skeletal Remains. J Forensic Sci. 2004;49(5): 1–5.

[46] Rennick SL, Fenton TW, Foran DR. The effects of skeletal preparation techniques on DNA from human and non-human bone. J Forensic Sci. 2005; 50(5):1016–1019.

[47] Lanting JN, van der Plicht J. Wat hebben Floris V, skelets Swifterbant S2 en visotters gemeen? Palaeohistoria. 1996;37/38: 491–520.

[48] Ervynck A, Boudin M, Van Neer W. Assessing the radiocarbon freshwater reservoir effect for a northwest-european river system (the schelde basin, Belgium). Radiocarbon 2018;60(2): 395–417

